# Dr *AFC*: Drug Repositioning Through Anti-Fibrosis Characteristic

**DOI:** 10.1101/2020.03.30.015123

**Authors:** Dingfeng Wu, Wenxing Gao, Xiaoyi Li, Chuan Tian, Na Jiao, Sa Fang, Jing Xiao, Zhifeng Xu, Lixin Zhu, Guoqing Zhang, Ruixin Zhu

## Abstract

Fibrosis is a key component in the pathogenic mechanism of many diseases. These diseases involving fibrosis may share common mechanisms, therapeutic targets and therefore, common intervention strategies and medicines may be applicable for these diseases. For this reason, deliberately introducing anti-fibrosis characteristics into modelling may lead to more success in drug repositioning. In this study, anti-fibrosis knowledge base was first built by collecting data from multiple resources. Both structural and biological profiles were derived from the knowledge base and used for constructing machine learning models including Structural Profile Prediction Model (SPPM) and Biological Profile Prediction Model (BPPM). Three external public data sets were employed for validation purpose and further exploration of potential repositioning drugs in wider chemical space. The resulting SPPM and BPPM models achieve area under the receiver operating characteristic curve (AUC) of 0.879 and 0.972 in the training set, and 0.814 and 0.874 in the testing set. Additionally, our results also demonstrate that substantial amount of multi-targeting natural products possess notable anti-fibrosis characteristics and might serve as encouraging candidates in fibrosis treatment and drug repositioning. To leverage our methodology and findings, we developed repositioning prediction platform, Drug Repositioning based on Anti-Fibrosis Characteristic (Dr *AFC*) that is freely accessible via https://www.biosino.org/drafc.

## Introduction

Fibrosis is defined as the process of excessive accumulation of fibrous connective tissue in most tissues or organs, where normal cells are replaced by the extracellular matrix (ECM), resulting in disrupted tissue function. In the new era of 21st century, the morbidity and mortality rates of various fibrotic diseases have increased progressively, bringing a huge global health burden. In developed countries, fibroproliferative diseases are responsible for nearly 45% of deaths[1]. One of the well-known fibrotic diseases, idiopathic pulmonary fibrosis(IPF), has a poor prognosis with the 5 year survival rate less than 30% and median survival ranging from 3 to 5 years[2]. The outcomes of IPF patients are even worse than those with many types of cancers [3]. As data obtained by Clinical Practice Research Datalink(CPRD) revealed, the prevalence of IPF patients in board case definitions has doubled from 19.94 per 100,000 patients in 2000 to 38.82 per 100,000 patients in 2012, and a 80% increase in incidence was observed[4]. Another life-threatening fibrotic disease, cardiac fibrosis, is one of the leading factors causing heart failure (HF) [5]. A research from 2008-2014 revealed that in 318 patients with systolic dysfunction, 78% had one type of myocardial fibrosis while 25% had at least 2 types [6].

The polypharmacology of most anti-fibrosis drugs could improve therapeutic efficacy. Recent studies have found that, firstly, fibrosis is the common pathogenic process in most diseases. For example, there are multiple common cellular processes between lung cancer and IPF, including inflammation, cell apoptosis and tissue infiltration [7]. Secondly, fibrosis-related processes have common mechanisms, targets and drugs [8, 9]. A multi-organ fibrosis research discovered a set of 90 common differentially expressed genes across lung, heart, liver and kidney. In the two most active gene networks generated by Ingenuity Pathway Analysis(IPA), these genes play a key role in connective tissue disorders and genetic, skeletal and muscular disorders[10]. Similarly, another multi-organ fibrosis research also obtained a series of 11 metzincin-related differentially expressed genes across heart, lung, liver, kidney and pancreas including *THBS2, TIMP1, COL1A2, COL3A1, HYOU1, MMP2* and *MMP7*[11]. Thirdly, fibrosis is a complicated pathological process involving multiple pathways, thus multi-target drugs are appropriate for fibrosis-related diseases[9]. Different pathways interact and counter-interact with each other to establish a “check-and-balance” system, for instance, the core regulators, transforming growth factor-β(TGF-β) and connective tissue growth factor(CTGF) signaling pathways could collaborate to elicit pulmonary and renal fibrosis[12, 13]. In summary, these evidences indicate that anti-fibrosis intervention strategies and medicines may be applicable for more diseases through targeting their common fibrosis-related mechanisms. Therefore, compounds that can more specifically target anti-fibrosis could have greater potential of repositioning and are more applicable for drug repositioning research.

Drug repositioning, or repurposing refers to the “reuse of old drugs”, recycling existing drugs for new medical indications. Compared with *de novo* drug discovery, drug repositioning has obvious advantages that it could significantly shorten drug development periods, reduce laboratory cost and minimize potential safety risk. Nowadays, drug repositioning is one of the most efficient strategies in drug development[14]. With the advancement of high-throughput sequencing technology and deep learning, various data-driven computational prediction and analytic models stand out[15, 16], including Similarity Ensemble Approach (SEA)[17] and Connectivity Map(cMAP)[18]. SEA clusters ligands into sets and calculates the similarity scores between ligand sets from ligand topology[17]. cMAP computes the similarity of “signatures” deduced from compound-induced gene profiles to quantify the biological functional relationships between compounds. Moreover, the relationship between compounds and diseases could also be quantified in opposite manner[18]. However, with so many repositioning methods and algorithms have emerged[19-21], there still no attempts hitherto in introducing anti-fibrosis characteristic into drug repositioning strategy.

For the first time, we built the anti-fibrosis knowledge base from anti-fibrosis related research. Based on the knowledge base, two repositioning models, Structural Profile Prediction Model (SPPM) and Biological Profile Prediction Model (BPPM) were constructed with high prediction accuracy. Centered on these two models, we then developed a repositioning computing platform, Drug Repositioning based on Anti-Fibrosis Characteristic (Dr *AFC*), to accelerate the process of exploring repositioning drugs and studying its underlying mechanisms.

## Materials and methods

### Datasets

#### Anti-fibrosis knowledge base

Anti-fibrosis related literatures were collected through key word queries “fibrosis AND target” in PubMed from Jan. 1st, 2000 to Oct. 31st, 2019. The compound-target interaction information on “fibrosis” were collected in the CTD[22] from Jan. 1st, 2000 to Oct. 31st, 2019. Anti-fibrosis trials were collected in ClinicalTrials.gov[23] from Jan. 1st, 2000 to Oct. 31st, 2019. Finally, anti-fibrosis treatments, targets and compound-target interactions were extracted and aggregated into the knowledge base.

#### Model construction

Structural and biological profiles of compounds were collected from DrugBank[24] and cMap, respectively and used for model construction. 2640 approved drugs in DrugBank and 1223 compounds in the anti-fibrosis knowledge base served as the raw data for Structural Profile Prediction Model (SPPM) construction. 6100 biological profiles (gene expression) of 1309 small molecules in cMap served as the raw data for Biological Profile Prediction Model (BPPM).

#### Case studies

20,263 natural products from TCMID[25], 5968 DrugBank experimental drugs[24] and 5000 random compounds from ChEMBL[26] were collected as external validations and case studies of SPPM. And external biological profiles from GEO database (GSE85871) that contains transcriptomics perturbation profiles of 105 natural products in MCF7 cell line were used for case studies of BPPM.

## Methods

### Pre-processing of modeling data

In raw chemical structures (from DrugBank approved drugs and the anti-fibrosis knowledge base) and biological profiles (from cMap) data, compounds that appeared in the anti-fibrosis knowledge base were labeled as positive candidates while the rest were labeled as negative candidates. Then, chemical structures were converted into chemical fingerprints (166-bits MACCS keys) for processing chemical information in a fast and convenient way using RDKit[27]. As to biological profiles, Quantile Transformer was used to transform biological profiles into ranking orders to improve the performance of model generalization, and also made datasets from different batches and platforms more comparable.

One-class SVM (nu=0.3) was performed to estimate sample quality, remove outliers and confirm final positive and negative samples. 70% of final samples were used as training set for model selection and super-parameter determination while the remainder as testing set for model validation.

### Anti-fibrosis model construction and validation

Four different machine learning algorithms were selected for modeling on training set, including logistic regression, decision tree, random forest and gradient boosting. Among them, method with highest precision and AUC calculated by 5-fold cross-validation was selected for subsequent analysis. Iterative feature elimination (IFE) algorithm was performed to select optimal feature set through one-by-one feature deletion. Finally, SPPM and BPPM were constructed based on optimal modeling algorithm and feature set, and further validated by testing set.

### Drug repositioning mechanism analysis

Network-based inference approaches were wildly used in drug repositioning [20, 21]. Here we infer the potential drug repositioning mechanism through compound-target-disease network. Firstly, based on SPPM and BPPM, the repositioning characteristics of compounds were predicted through their structural or biological profiles, in which compounds with reposition score>0.5 were considered as anti-fibrosis and had repositioning potential. Next, the anti-fibrosis characteristic and potential repositioning mechanisms of these candidates were explored on the basis of compound-target-disease corresponding information in the anti-fibrosis knowledge base. Similar compounds that may interact with same targets and diseases were calculated through Tanimoto similarity of chemical structural fingerprints or Spearman’s rank correlation coefficient of biological profiles. Targets and disease information of compounds reported in previous researches were refined from the anti-fibrosis knowledge base to explore anti-fibrosis mechanism of compounds. Finally, the potential mechanisms among compounds in compound-target-disease network displayed in drug repositioning analysis were used to help propose feasible drug repositioning solutions.

### Webserver construction of Dr AFC

Dr *AFC* was constructed through PostgreSql database and Django framework. This platform serves as a practical tool for prediction of drug repositioning potential based on compound structures (SPPM) and biological profiles (BPPM) as well as displaying compound-target-disease network of drug repositioning mechanisms. Meanwhile, Dr *AFC* also integrated toolkits such as quantitative estimate of drug-likeness (QED) from Silicos-it[28], and similarity calculation and structure matching borrowed from RDkit to provide convenient web-based calculations for users.

The overall process is shown in Figure 1.

**Figure 1.**
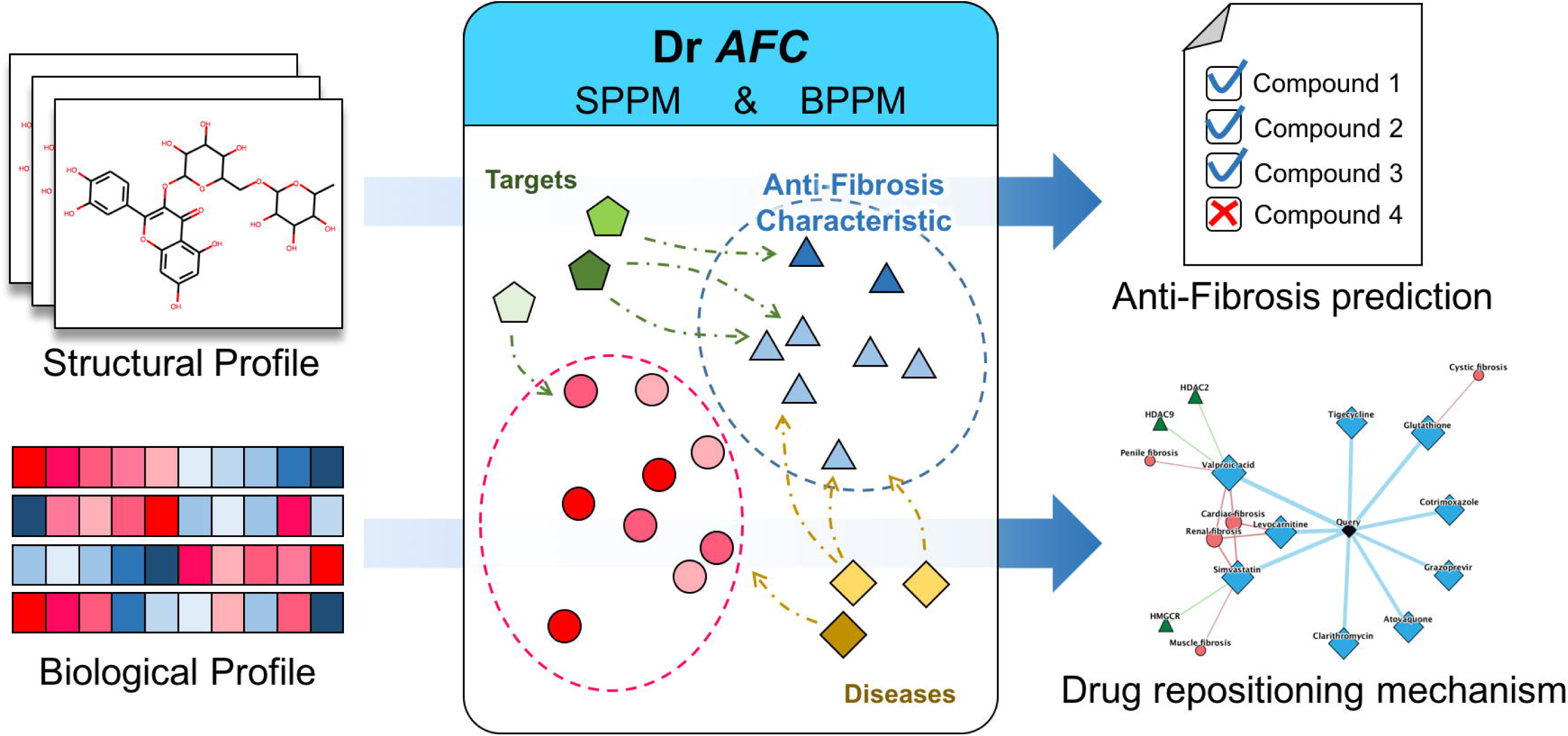
The schematic of Dr *AFC* construction.

## Results

### SPPM and BPPM show high performances for anti-fibrosis prediction

To construct the anti-fibrosis knowledge base, 7058 fibrosis-related references from PubMed, 302 from Comparative Toxicogenomics Database(CTD)[22] and 2664 fibrosis-related trials from ClinicalTrials.gov[23] were collected through text mining. Finally, 1223 anti-fibrosis treatments (containing 902 small molecules), 1067 fibrosis-related targets, 3096 fibrosis-related records from references and 1787 from trials, 1067 anti-fibrosis compound-target interactions were obtained and integrated into anti-fibrosis knowledge base (Figure S1).

In modeling session, 2885 compound structures (from DrugBank approved drugs) [24] and 6100 biological profiles (from cMap) were labeled as positive candidates and negative candidates based on their anti-fibrosis characteristic in the anti-fibrosis knowledge base. After sanity check and outlier removal, 1701 compound structures and 2735 biological profiles were filtered out for model construction (Table S1).

Four different machine learning classifiers were evaluated and compared to choose the most optimal modeling method (Table S2). Gradient boosting was eventually selected according to its highest precision and AUC (Structural profile: Precision=0.737, AUC=0.839, Biological profile: Precision=0.892, AUC=0.912).

In the process of building SPPM and BPPM, we found that even a small number of features could reach certain stability and reasonably good performance (Figure S2, **Figure 2a**). Models based on top 38 features including CHARGE, S and XA(A)A could reach the maximum cross-validation AUC (0.879) in SPPM while top 47 features including RPL30, MRMRPL5 and KPNB1 could reach the maximum cross-validation AUC(0. 972) in BPPM. We discovered that 46 of the top 47 features in BPPM were connected with fibrosis in CTD inference networks (Figure 2b). Besides, several genes were associated with fibrosis-related indications like retroperitoneal fibrosis, keloids, tissue adhesions and cicatrix.

**Figure 2.**
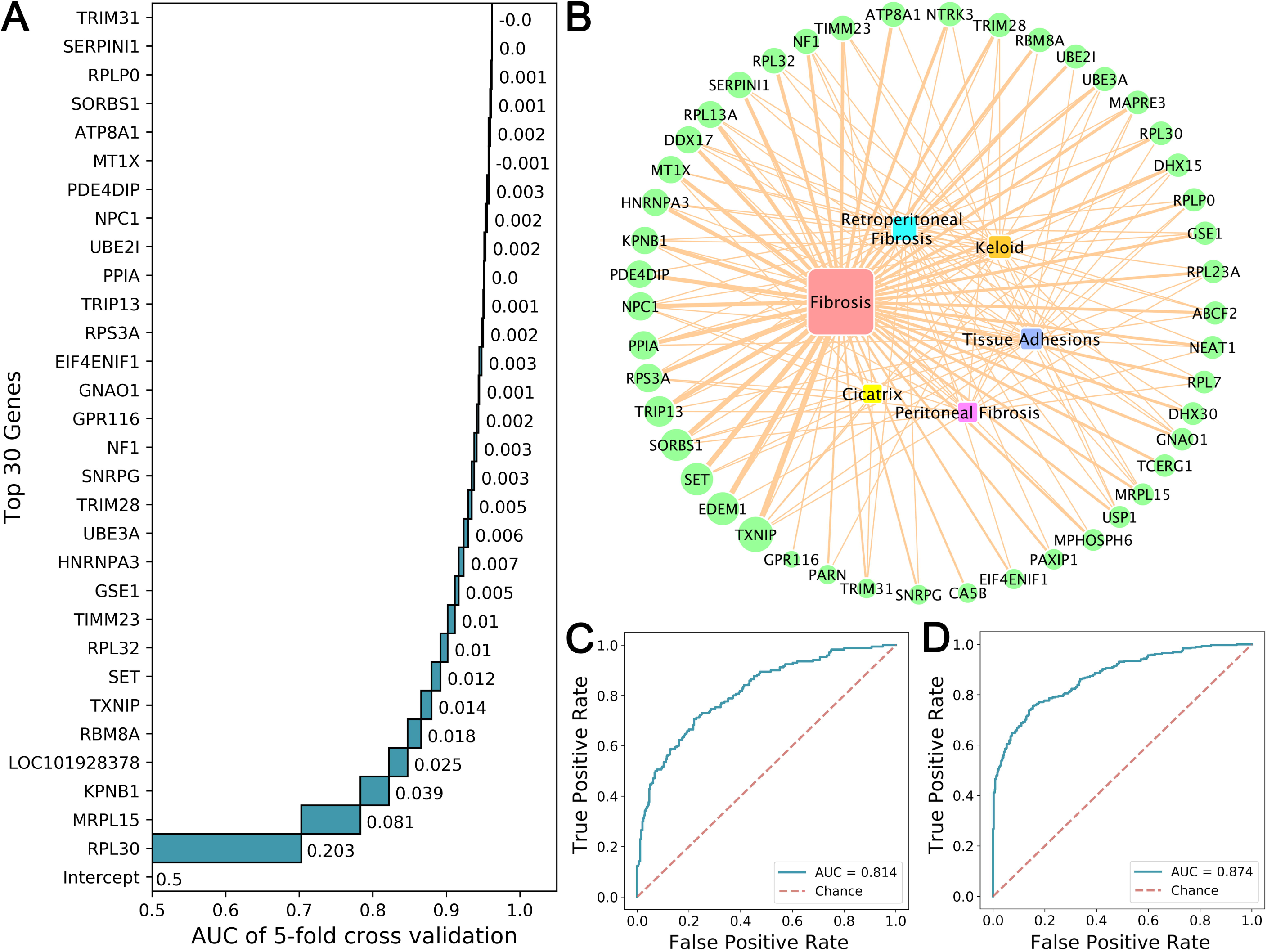
Feature selection and model performances. **A**. Performances of top 30 features through iterative feature elimination in BPPM; **B**. The CTD inference networks of 47 gene features and fibrosis-related diseases; **C**. AUC of SPMM in testing set; **D**. AUC of BPPM in testing set.

Finally, SPPM and BPPM were build based on the most optimal modeling method and the selected small feature subset (top 38 features in SPPM and top 47 features in BPPM). In testing set, the average AUC for SPMM reaches 0.814 (Figure 2c) while the average AUC for BPMM reaches 0.874 (Figure 2d).

### Case studies

#### Anti-fibrosis drugs exhibit greater drug repositioning potential

We used SPPM to predict anti-fibrosis drugs from DrugBank experimental drugs and the comparative analysis was performed between the CTD compound-gene interactions of the predicted anti-fibrosis and non-anti-fibrosis drugs. The results show that the anti-fibrosis group accommodates stronger interactions, presumably more genetic effects thus greater repositioning potential (**Figure 3a**).

**Figure 3.**
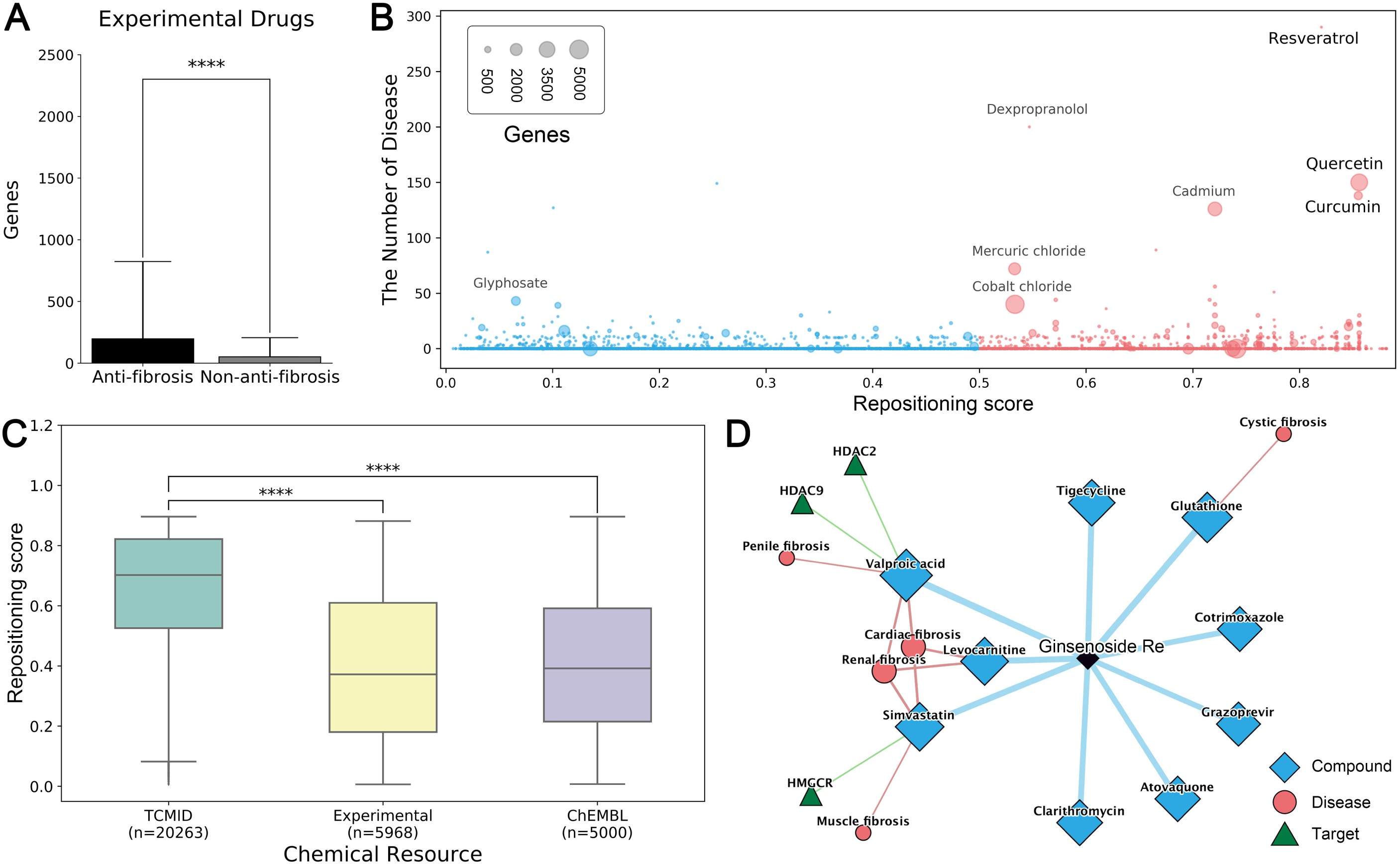
Case studies of Dr *AFC*. **A**. Comparison of the number of genes interacting with compounds predicted as anti-fibrosis and non-anti-fibrosis; **B**. The distribution of related genes, diseases and repositioning score for Drugbank experimental drugs. Compounds with repositioning score>0.5 were considered as anti-fibrosis and had repositioning potential; **C**. The distribution of repositioning scores in different datasets (****: p-value <10^−4^ by two-sided Wilcoxon rank sum test); **D**. Drug repositioning mechanism analysis of ginsenoside Re by Dr *AFC*.

In Drugbank experimental drugs, multiple drugs with great repositioning potential (Related Genes>500 and Diseases>20, Figure 3b, Table S3) were developed for fibrotic diseases and other diseases. Quercetin was discovered to ameliorate liver fibrosis through regulating macrophage infiltration and polarization, and it could alleviate IPF through fibroblasts apoptosis[29, 30]. Based on our results, we confirm that quercetin interacts with numerous genes and is strongly linked to multiple diseases (Repositioning score=0.856, Related Genes=3938, Diseases=150, Table S3). Another natural compound from turmeric, curcumin (Repositioning score=0.855, Related Genes=903, Diseases=138, Table S3), could also be used for treating multiple fibrotic diseases. It could inhibit fibroblast proliferation and myofibroblast differentiation in IPF[31] while inhibit oxidative stress and exhibit anti-inflammatory effect in liver fibrosis[32]. Apart from fibrosis, curcumin has been applied for osteoarthritis and rheumatoid arthritis treatment [33, 34]. Moreover, other drugs, such as resveratrol also had great repositioning potential (Repositioning score=0.821, Figure 3b).

#### Natural compounds are the better repositories for drug repositioning

In order to expand the resources of potential repositioning drugs and further explore the chemical space, we introduced two external molecule sets, natural products from TCMID[25] and random compounds in ChEMBL[26]. SPPM was used to predict the repositioning potential of compounds from both external molecule sets. The results show that there were 35.42%, 77.26% and 37.04% of compounds could be potentially repositioned in DrugBank experimental drugs, TCMID and ChEMBL, respectively. The reserves in natural products from TCMID are significantly higher than others, indicating that natural products are great repositioning repositories and need further researches (Figure 3c).

BPPM was used to discover specific natural products with repositioning potential from gene profiles dataset of 105 natural products (GSE85871). The results show that a total of 66 natural products have anti-fibrosis characteristic and repositioning potential, including ginsenoside Re(Repositioning score=0.979), muscone(Repositioning score=0.974) and cinnamic acid(Repositioning score=0.948) (Table S4). Among them, ginsenoside Re hold the potential to influence HDAC2, HDAC9 and HMGCR and fulfilled anti-fibrosis roles via “inflammation”, “preventing collagen deposition” and “targeting myeloperoxidase” with Drug repositioning mechanism analysis tools in Dr *AFC* (Figure 3d). Ginsenoside Re is the extract of panax ginseng which exhibited protective effects in neural and systematic inflammations through inhibiting the interaction between LPS and TLR4 in macrophages[35]. It was reported to exert anti-fibrosis effect on cardiac fibrosis through down-regulating the expression of p-Smad3, collagen I and reducing the augmentation of collagen fibers[36]. Apart from fibrosis, ginsenoside Re could alleviate inflammation through inhibiting myeloperoxidase activity[37] and decrease fat accumulation through inhibiting HMGCR and cholesterol biosynthesis[38]. Besides, other ginsenosides, like ginsenoside Rb1, ginsenoside Rc, ginsenoside Rb3, ginsenoside Rb2, ginsenoside Rd and ginsenoside Rg, also exhibit anti-fibrosis characteristic and repositioning potential(Table S4).

### Drug Repositioning based on Anti-Fibrosis Characteristic Webserver

Based on SPPM and BPPM, we constructed a computing platform for repositioning research purpose, named Drug Repositioning based on Anti-Fibrosis Characteristic (Dr *AFC*), the main function and workflow of which is shown in **Figure 4**. On Dr *AFC* platform, anti-fibrosis and potential repositioning could be predicted from compound structures or biological profiles. Drug repositioning mechanism analysis could infer the relationships among compounds, fibrosis-related targets and diseases which help understand pathology. Furthermore, drug-likeness estimation, chemical similarity calculation and structure matching were integrated into Dr *AFC* to provide useful information for drug development.

**Figure 4.**
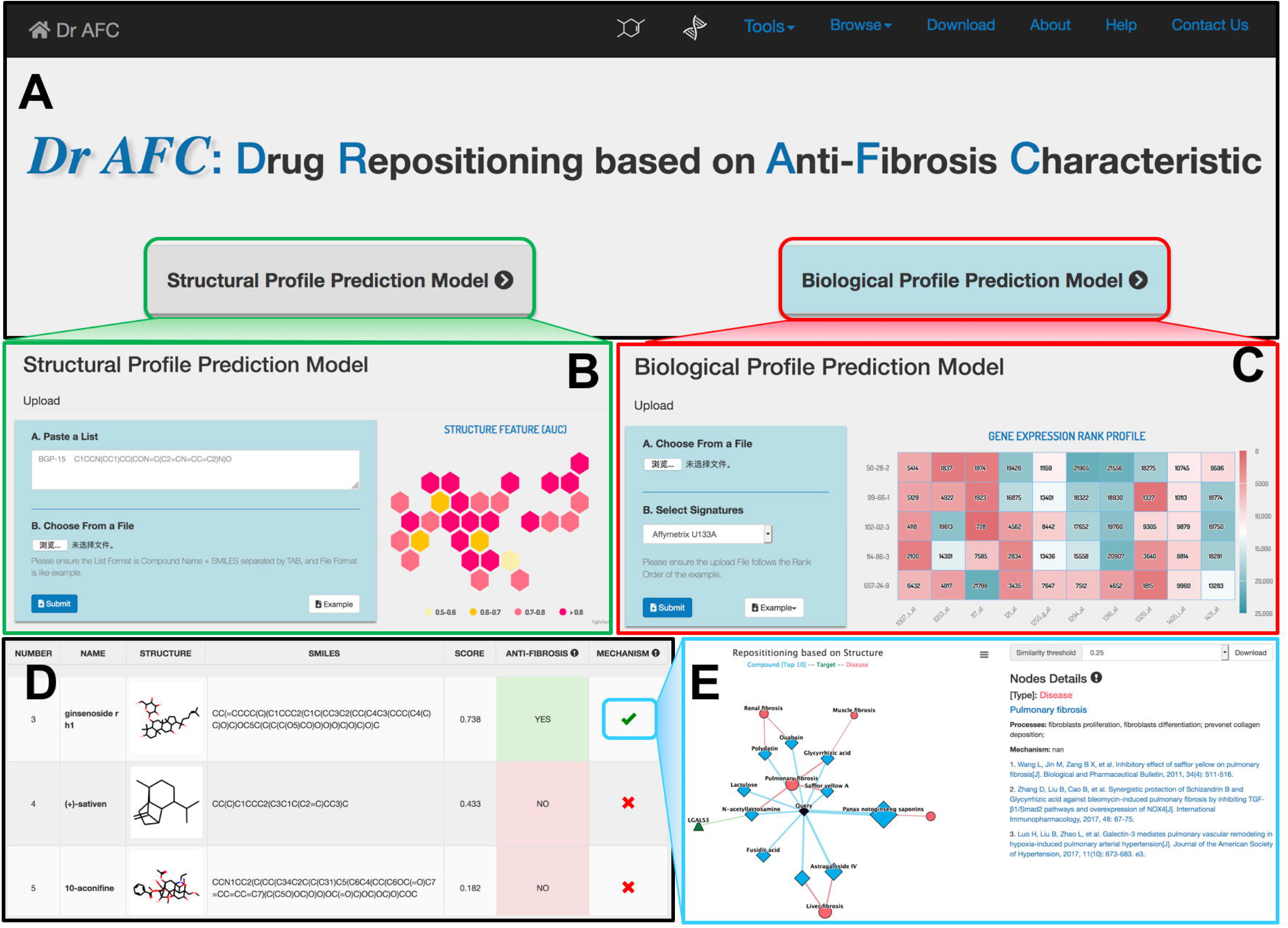
Anti-fibrosis and repositioning computing platform (Dr *AFC*) **A**. Dr *AFC* integrated two prediction models, SPPM and BPPM; **B**. SPPM accepts SMILES strings of chemical structures in text or file; **C**. BPPM accepts biological profiles in file; **D**. Repositioning score, label and functional network of compounds were displayed in result; **E**. Drug repositioning mechanism analysis was implemented to infer the drug potential repositioning mechanism through relationships among similar compounds, fibrosis-related targets and diseases.

#### Drug repositioning analysis function

Dr *AFC* allows users to upload compound structures or compound-induced biological profiles for repositioning potential prediction. As shown in Figure 4b, Dr *AFC* accepts SMILES strings of compound structures for SPPM prediction, and accepts gene profiles with row names in Affymetrix U133A probe ID, Entrez ID or gene symbol format for BPPM prediction. Both methods support .txt,.csv or .xlsx files (Figure 4c).

Webserver would perform corresponding prediction analysis automatically based on the uploaded files and display the output on the result page in three aspects (Figure 4d): 1) Basic part includes compound ID, compound name, 2D compound structure(only for SPPM) and SMILES string(only for SPPM). 2) Prediction part includes repositioning scores of anti-fibrosis characteristic and repositioning potential prediction. The repositioning scores ranges from 0 to 1 and higher score indicates higher potential. If repositioning score≥0.5, the compound would be defined as an anti-fibrosis and potential repositioning compound. 3) Drug repositioning mechanism analysis part. This analysis infers the potential anti-fibrosis and repositioning mechanisms of compound structures or biological profiles users uploaded based on our anti-fibrosis knowledge base. It could provide users potential mechanisms as theoretical foundations for drug repositioning studies.

#### Other functions

Dr *AFC* also contains drug-likeness estimation, chemical similarity calculation and structure matching tools. Users could upload their compounds in SMILES and perform these additional functions. Drug-likeness estimation could evaluate and score the compound drug-likeness, which ranges from 0 to 1 with higher score indicating higher potential for lead compound. Chemical similarity calculation and structure matching provide convenient ways for users to search compound with similar structures, same structures or substructures, supporting single compound calculation and simultaneous calculation for multiple compounds.

## Discussion

Fibrosis is the common mechanism of diseases that attracts global attention. The anti-fibrosis characteristic of a compound could infer the greater repositioning potential it would have. However, the anti-fibrosis characteristic has not been extensively introduced into the realm of drug discovery till now. In this study, we first bridge the gap by developing a platform that can provide intensive information conveniently on drug repositioning based on anti-fibrosis characteristic data, Dr *AFC* (https://www.biosino.org/drafc). This *in silico* platform also provides a highly accurate way to generate data for rational drug design via combining the advanced machine-learning algorithm.

Dr *AFC* was built based on the anti-fibrosis knowledge base, which pioneered the excavation and organization of fibrosis-related studies throughout recent years. Structural profile (SPPM) and biological profile (BPPM) that show extraordinary capabilities in drug repositioning prediction (with AUC 0.814 and 0.874, respectively) were integrated into Dr *AFC*. BPPM show slightly higher performance than SPPM according to the AUC. The possible reason could be that biological profile is more tolerant and could contain information reflecting an overall effect of compound functionally in the body. Biological profile show its advantage in multiple repositioning algorithms previously, such as cMAP[18], L1000CDS^2^[39] and MANTRA[40]. Besides, certain therapies without available structure profile like biotech drugs or cocktail therapies could also be studied in repositioning research according to their biological profiles.

In BPPM, 47 biological markers exhibited strong prediction abilities. These genes are directly or indirectly linked to various fibrotic diseases. Interestingly, ribosomal proteins including RPL30, MRPL15, RPL32, RPS3A, RPLP0, RPL7, RPL23A and RPL13A are the main part of these biological markers. Ribosomes serve as significant regulators in immune signaling pathways, tumorigenesis pathways and cardiovascular and metabolic diseases[41, 42]. For example, the expression of RPL30 is negatively correlated with carcinogenesis process in medulloblastoma that usually is accompanied by desmoplasia and could thus serve as a prognosis biomarker[43]. Besides, the over-activation of RNA polymerase in the biogenesis of ribosomes could cause the enhancement of protein synthesis and the decrease of translation accuracy, triggering cancers or exacerbating cancer processes[44]. Furthermore, some biological markers are associated with the spliceosome formation including RBM8A, HNRNPA3, SNRPG and DHX15. Spliceosome is the large molecular machine composed of five snRNA and many proteins, and serves as the catalyzer of pre-RNA introns which are crucial for protein expression and function. It has been reported to be closely associated with multiple diseases, including cystic fibrosis and pulmonary fibrosis[45, 46].

Based on external molecule sets, natural products are validated to have the strongest anti-fibrosis characteristics and repositioning potential among chemicals from different sources. Natural products provide a wealth of valuable natural resources for modern medicine and are seen as promising and popular candidates for drug repositioning studies[47]. Their privileged scaffolds, structural complexity, abundant stereochemistry and ‘metabolite-likeness’ are main reasons for the broad-spectrum of biological activities [48, 49]. The multi-targets and synergistic effects of natural products exhibit great advantages in treating diseases undergoing sophisticated mechanisms, such as fibrosis[50]. Our studies show that natural products like ginsenoside have great anti-fibrosis characteristic and repositioning potential and should be top priority when considering repositioned drug discovery. Additionally, the natural products in Drugbank experimental drugs such as quercetin, curcumin and resveratrol, also highlight their strong repositioning capabilities. Therefore, natural products could serve as promising source and the good choice for further drug development and repositioning study.

## Conclusion

In summary, based on anti-fibrosis characteristics, we constructed two repositioning models, SPPM and BPPM, which could predict the anti-fibrosis characteristics and repositioning potential from compound structures and compound-induced biological profiles. SPPM and BPPM efficiently utilize the generality of fibrotic diseases, thus greatly increase the success rate of drug repositioning. This study not only established a highly efficient strategy of prediction, but also developed a convenient and user-friendly computing platform, Dr *AFC* (https://www.biosino.org/drafc), for studying fibrosis mechanisms and drug repositioning.

## Key Points

- Fibrosis is the common mechanism of diseases which could be applied in drug repositioning.
- We developed a convenient and user-friendly computing platform, Dr *AFC*, for studying fibrosis mechanisms and drug repositioning.
- Dr *AFC* shows high performance on both cross validation and external validation, which demonstrates its potential applications in drug discovery.
- Natural compounds proved to be the better repositories for drug repositioning.

## Supplementary material

**Figure S1 Knowledgebase architecture of Dr AFC**

**Figure S2 Performances of top 30 features through iterative feature elimination in SPPM**

**Table S1 The sample size of SPPM and BPPM**

**Table S2 Performances of four different machine learning classifiers**

**Table S3 Drug repositioning prediction in Drugbank Experimental drugs**

**Table S4 Drug repositioning prediction in natural products(GSE85871)**

