## Supplementary figures and images for "Dr *AFC*: Drug Repositioning Through Anti-Fibrosis Characteristic"

### Figure S1

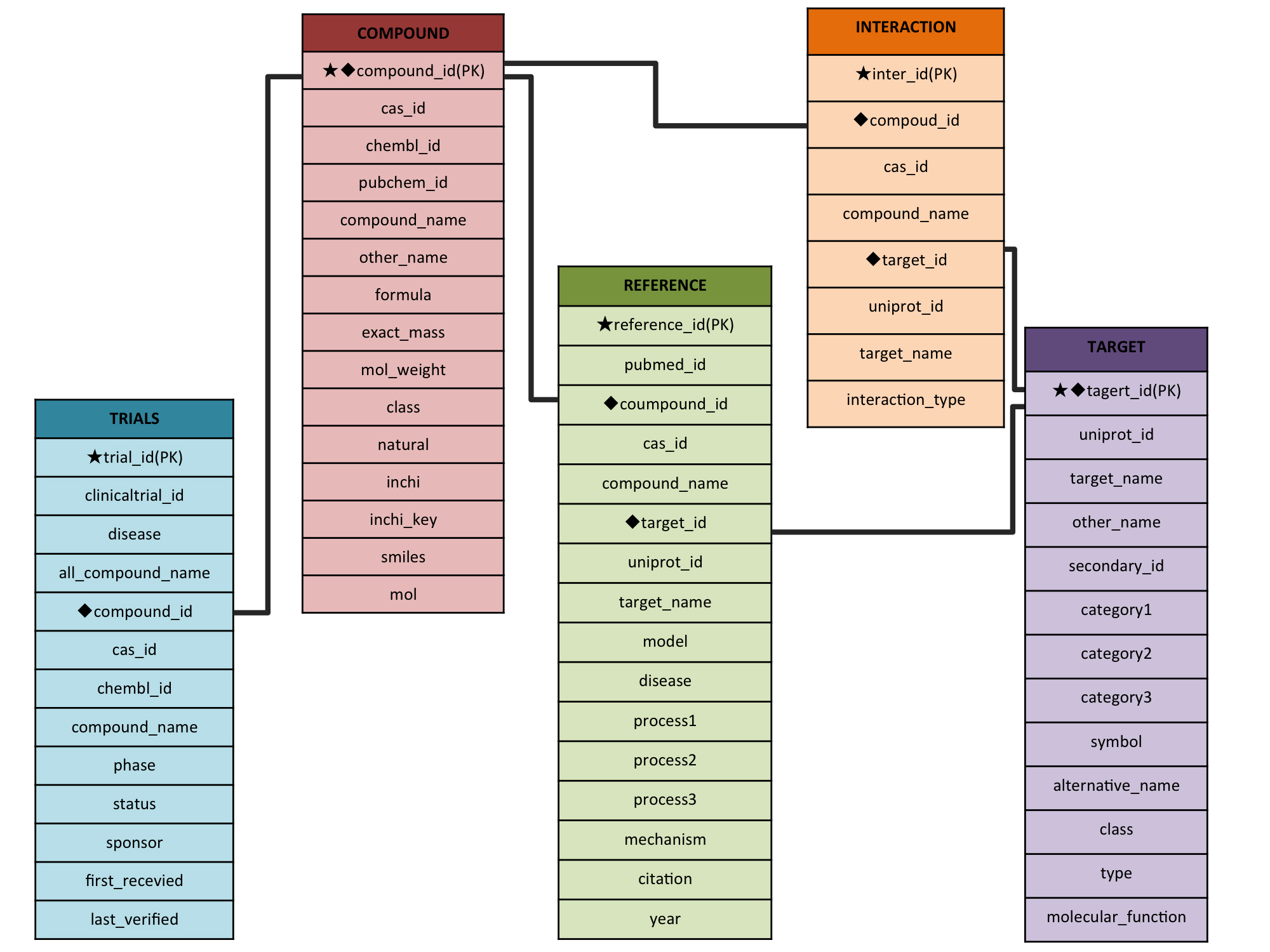

### Figure S2

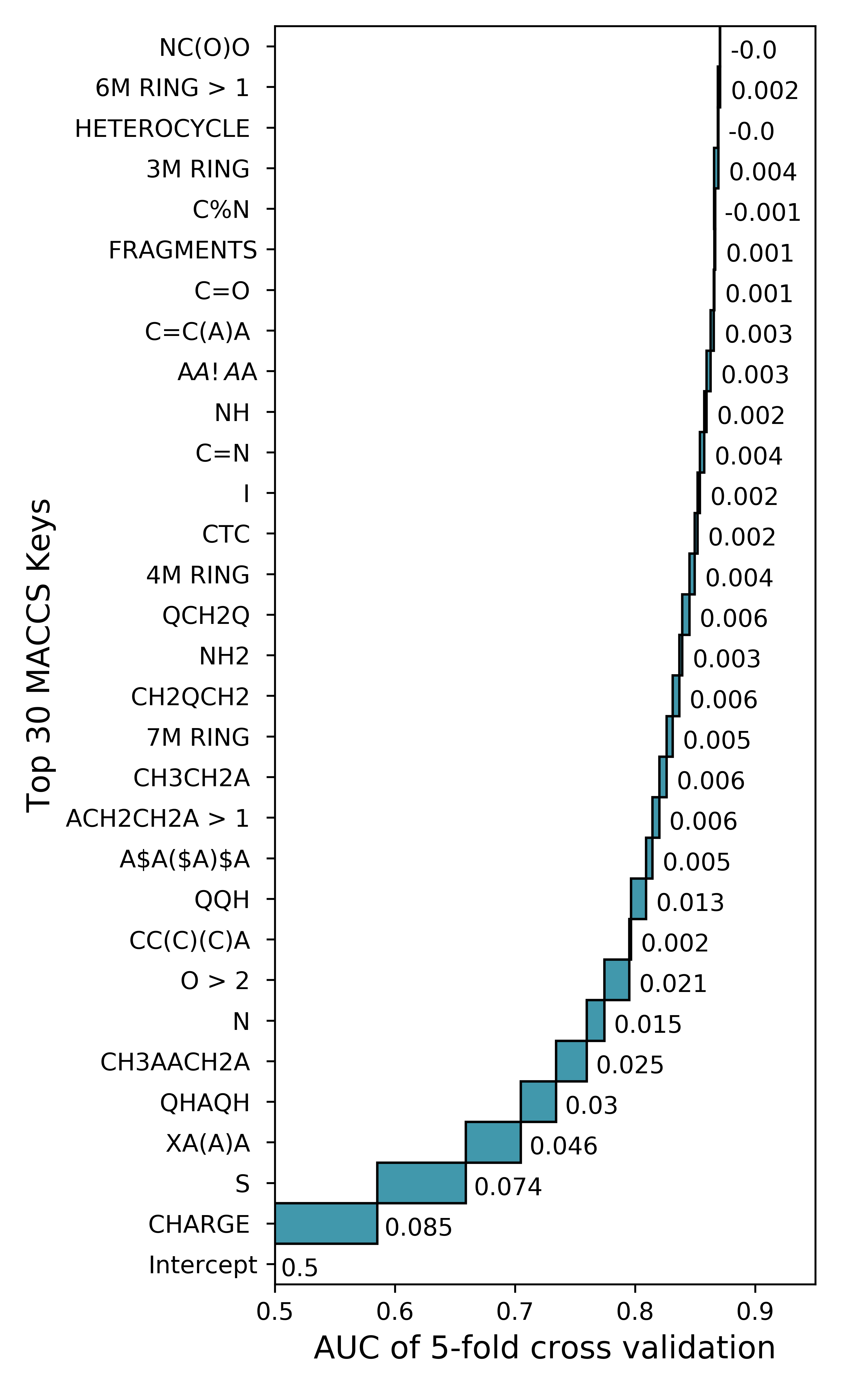
